# Exploratory analyses suggest less cognitive decline with nilvadipine treatment in very mild Alzheimer’s disease subjects

**DOI:** 10.1101/345652

**Authors:** L Abdullah, F Crawford, H Langlois, M Tsolaki, A Börjesson-Hanson, M Olde Rikkert, F Pasquier, A Wallin, S Kennelly, S Hendrix, K Blennow, B Lawlor, M Mullan

## Abstract

**Background:** We explored whether the effects of nilvadipine on cognition were influenced by baseline Alzheimer’s disease (AD) severity.

**Methods:** Exploratory analyses were performed on the modified intention-to-treat (mITT) dataset (n = 497) of a phase III randomized placebo-controlled trial to examine the response to nilvadipine in very mild, mild and moderate AD subjects. The outcome measures included total and subscale scores of the Alzheimer’s Disease Assessment Scale Cognitive 12 (ADAS-Cog 12), the Clinical Dementia Rating Scale sum of boxes (CDR-sb) and the AD composite score (ADCOMS), an outcome measure recently developed to detect treatment responses in subjects with prodromal AD. Cerebrospinal fluid (CSF) biomarkers Aβ38, Aβ40, Aβ42, total tau and P181 tau were measured in a subset of samples (n = 55). Regression analyses were adjusted for potential confounders and effect modifiers in order to examine the interactive effects of nilvadipine and AD severity on cognitive outcomes over 78-weeks.

**Results:** Compared to their respective placebo-controls, nilvadipine-treated, very mild AD subjects showed less decline, whereas moderate AD subjects showed greater decline on the ADAS-Cog 12. Also in very mild AD, a beneficial effect (as measured by ADCOMS), was detected in the nilvadipine treated group. Therapeutic effects of nilvadipine were also observed for a composite memory trait in very mild AD subjects and a composite language trait in mild AD subjects. CSF Aβ42/Aβ40 ratios were increased in mild AD and decreased in moderate AD patients treated with nilvadipine, compared to their respective controls.

**Conclusion:** These data suggest that very mild AD subjects benefited from nilvadipine and that future clinical trials of nilvadipine in this population are required to confirm these findings.

**Trial Registration:** NCT02017340 Registered 20 December 2013, https://clinicaltrials.gov/ct2/show/NCT02017340

EUDRACT Reference Number 2012-002764-27 Registered 04 February 2013, https://www.clinicaltrialsregister.eu/ctr-search/search?query=2012-002764-27

## Introduction

Alzheimer’s disease (AD) is the most common neurodegenerative disease, affecting nearly 5.3 million US citizens. By 2050, the prevalence of AD is expected to reach 13 million in the US alone and 100 million worldwide. The presence of amyloid plaques and neurofibrillary tangles in the brain are key hallmarks of AD (1-3) and are also accompanied by cerebrovascular disease, α-synuclein and TDP-43 deposits and inflammation (4-6). Recent clinical trials have shown that moderate AD patients, with established brain amyloid and tau pathologies, are unresponsive to anti-amyloid therapeutic approaches, although some trials have shown potential benefits in mild and early stage AD patients (7-11). As such, early and mild AD patient populations may be more appropriate for anti-amyloid and anti-tau approaches.

Findings of the Systolic Hypertension in Europe (SYST-EUR) trial of 2400 participants showed that intervention with nitrendipine, a dihydropyridine (DHP) calcium channel blocker (CCB) similar in structure to nilvadipine, resulted in a reduction of AD incidence (12). A small clinical trial of nilvadipine in mild cognitive impairment (MCI) patients showed reduced conversion to AD in the subjects treated with nilvadipine compared to those on amlodipine, which, in contrast to nilvadipine, does not penetrate the blood brain barrier (BBB) (13). Preclinical studies in mouse models of AD have shown that nilvadipine improves cognitive function, lowers Aβ production, increases Aβ clearance across the BBB and reduces tau hyper-phosphorylation and inflammation (14-17). *In vitro* screening of over 1000 DHPs, using cell-based assays, showed that inhibition of Aβ production is not a class effect of DHPs, as only a minority of them, such as nilvadipine and nitrendipine, demonstrated inhibitory action while others had no effect or even enhanced Aβ production (15, 18). Nilvadipine is a racemic compound and an examination of the individual isomers showed both (+)-and (-)-nilvadipine forms to have beneficial effects on amyloid and tau pathologies in an AD mouse model (17) while only the (+)-nilvadipine has the blood-pressure-lowering effects through antagonism of calcium channels. These studies suggest that the preclinical anti-amyloid and anti-tau effects of nilvadipine are not related to its CCB properties. It has been shown that nilvadipine inhibits spleen tyrosine kinase (Syk), which consequently downregulates amyloid production, tau phosphorylation and inflammation (17). As such, nilvadipine may represent a novel, multimodal, disease-modifying therapy for AD. To evaluate this, a phase III multi-center, double-blinded, randomized, placebo-controlled clinical trial was conducted in Europe to test its efficacy in treating AD (the NILVAD trial).

In the NILVAD trial, when analyzed as a single population according to the protocol, combined mild and moderate AD subjects did not benefit from nilvadipine treatment. However, subgroup analyses indicated that, compared to placebo-treated controls, nilvadipine-treated mild AD subjects (baseline MMSE ≥ 20) showed cognitive benefits whereas moderate AD subjects (baseline MMSE < 20) showed worsening of cognition (19). These findings support further exploration of the treatment effects of nilvadipine in AD patients at the earlier stages of the disease. Given the proposed amyloid and tau lowering mechanisms-of-action of nilvadipine (15, 17) and findings from other drug studies targeting the same (7), we hypothesized that the most pronounced beneficial effects of nilvadipine treatment would occur in the subjects who had enrolled in the study when they were in the earliest clinical stages of disease progression. We therefore further stratified the study population by AD severity at baseline in mild AD subjects and conducted unplanned, data-dependent, exploratory analyses of the existing dataset from the NILVAD trial (20). We anticipated that these exploratory analyses would help identify responsive subgroups of patients for recruitment into future clinical trials of nilvadipine or related compounds. In a subset of the study population for whom cerebrospinal fluid (CSF) samples were available, we examined whether CSF Aβ and tau changes differed between nilvadipine and placebo groups when stratified by baseline severity of AD.

## Methods

### Study design and participants

This 18-month phase III double-blind, placebo-controlled, randomized clinical trial was conducted in 9 countries across Europe (see elsewhere for additional details (21)) and funded by the European Commission under a Framework 7 Programme Health Theme collaborative project grant. A separate Scientific Advisory Board, an independent Ethics Advisory Board and an independent Data Safety Monitoring Board were involved in the oversight of the trial. The study protocol and associated documents were approved by Research Ethnics Committee and Institutional Review Boards (IRB) for all study sites (see the ethics statement below for the full list of IRB by each country). A written consent for trial participation was obtained following a full explanation of the risks and benefits of the trial to potential participants (see elsewhere for details on the study leaflets (21)). A written consent was obtained from patients who had the ability to provide a consent as well as from the caregivers at the screening visit prior to initiating the study process. The procedure for obtaining informed consent from a participant with reduced decision-making capacity was conducted in accordance to the national laws of each country and assessed by the relevant bodies in each country. The sample size calculations for the main trial were based on the mean difference of 3.5 and SD of 9 between the treated and control groups and previously described elsewhere (21). The block randomization was performed using an online system hosted by the Clinical Trial Unit at the King’s College London. Blocks of varying sizes were used. The randomization was at the subject level and stratified by country site, see elsewhere for more details (21). All study investigators and patients were blinded to the treatment assignment. There were no interim analyses in this trial.

Inclusion criteria for the study required that participants should be over the age of 50 and have a diagnosis of mild or moderate probable AD according to the established guidelines from the National Institute of Neurological and Communicative Disorders and Stroke/Alzheimer’s Disease and Related Disorders Association Inc. (NINCDS-ADRDA) and the Alzheimer’s Association, and having a baseline Mini-Mental State Examination (MMSE) score of >12 and ≤ 27 (21). A total of 569 subjects were screened for eligibility and 511 were randomized into the trial with 253 assigned to the nilvadipine group (one dropped out due to blood pressure measurements being out of range) and 258 were assigned to placebo. Of these 510 subjects, 11 were lost to follow-up and 2 withdrew consent, leaving 497 subjects in the modified intention-to-treat (mITT) dataset (see Figure 1). Data from subjects in the mITT set were used for these additional exploratory analyses below. At baseline, each subject was randomly assigned to either 8mg of Nilvadipine or placebo once a day, and each study subject was required to take the capsule orally after breakfast for 78 weeks. The primary outcome measures were the 12-item Alzheimer’s Disease Assessment Scale–cognitive sub-scale 12 (ADAS-Cog 12) and the Clinical Dementia Rating scale sum of boxes (CDR-sb), and these tests were administered at four time-points (baseline, and weeks 13, 52 and 78).

**Figure 1:**
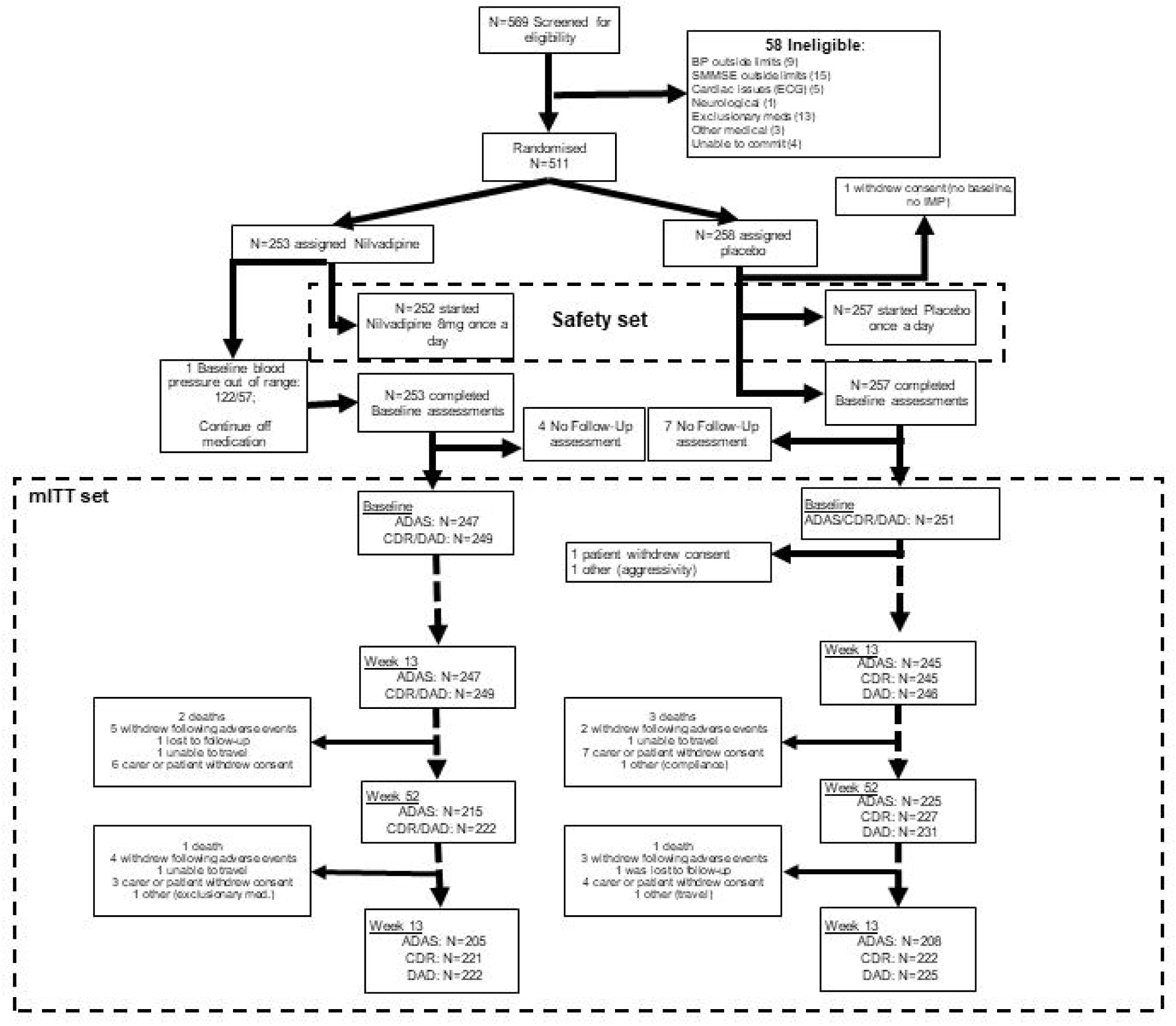
CONSORT flow-chart: Flow-chart of the main study from which data for exploratory analyses were obtained.

### Exploratory analyses

The exploratory analyses of the NILVAD trial were restricted to the co-primary outcome measures of ADAS-Cog 12 and CDR-sb. The mild AD group was further stratified by single point increases in the baseline MMSE scores ranging from 20 and above to 25 and above. From this data-driven subject stratification approach, we identified three AD severity subgroups as follows: moderate AD (baseline MMSE scores of ≤ 19), mild AD (baseline MMSE scores from 20-24) and very mild AD (baseline MMSE scores ≥ 25). The cutoff of MMSE score for the moderate AD category was chosen as previously recommended (20, 22). The nomenclature of mild and very mild AD was adopted in accordance with Huntley and colleagues (23). These analyses also explored the potential impact of nilvadipine treatment on cognitive sub-scales of the ADAS-Cog 12 and CDR-sb tests. The ADAS-Cog 12 sub-scales are: immediate word recall, delayed recall, naming, following commands, constructional praxis, ideational praxis, orientation, word recognition, remembering test directions and instructions, spoken language, comprehension and word finding difficulty in spontaneous speech. The sub-scales of CDR-sb are: memory, orientation, judgment and problem solving, community affairs, home and hobbies and personal care. In addition, ADAS-Cog 12 sub-scales were further grouped into specific traits for memory, language and praxis based on the topography of tissue loss in AD depending on the stage of the disease, as previously suggested by Verma and colleagues (24). Using this strategy, sub-scales related to each trait were grouped together to generate a composite variable for each trait (Figure 2A). We also calculated a modified AD Composite Score (ADCOMS) using the partial least square (PLS) coefficients previously generated by Wang and colleagues (25, 26). Briefly, PLS coefficients for each subscale from the ADAS-Cog 12 and CDR-sb were directly multiplied by scores for each subscale in our dataset. The ADAS-Cog 12 subscales were: delayed word recall, orientation, word recognition and word finding difficulty; the CDR-sb subscales were: personal care, community affairs, home and hobbies, judgment and problem solving, memory and orientation. Since we did not have MMSE scores at follow-up visits, the MMSE orientation was substituted with the ADAS-Cog 12 orientation subscale by dividing by 8 and multiplying by 5 so that it was on the same scale as the MMSE orientation score. Since the MMSE item constructional praxis accounts for only 0.04 points out of 1.97 of the PLS weight, this item was omitted from the analyses.

**Figure 2:**
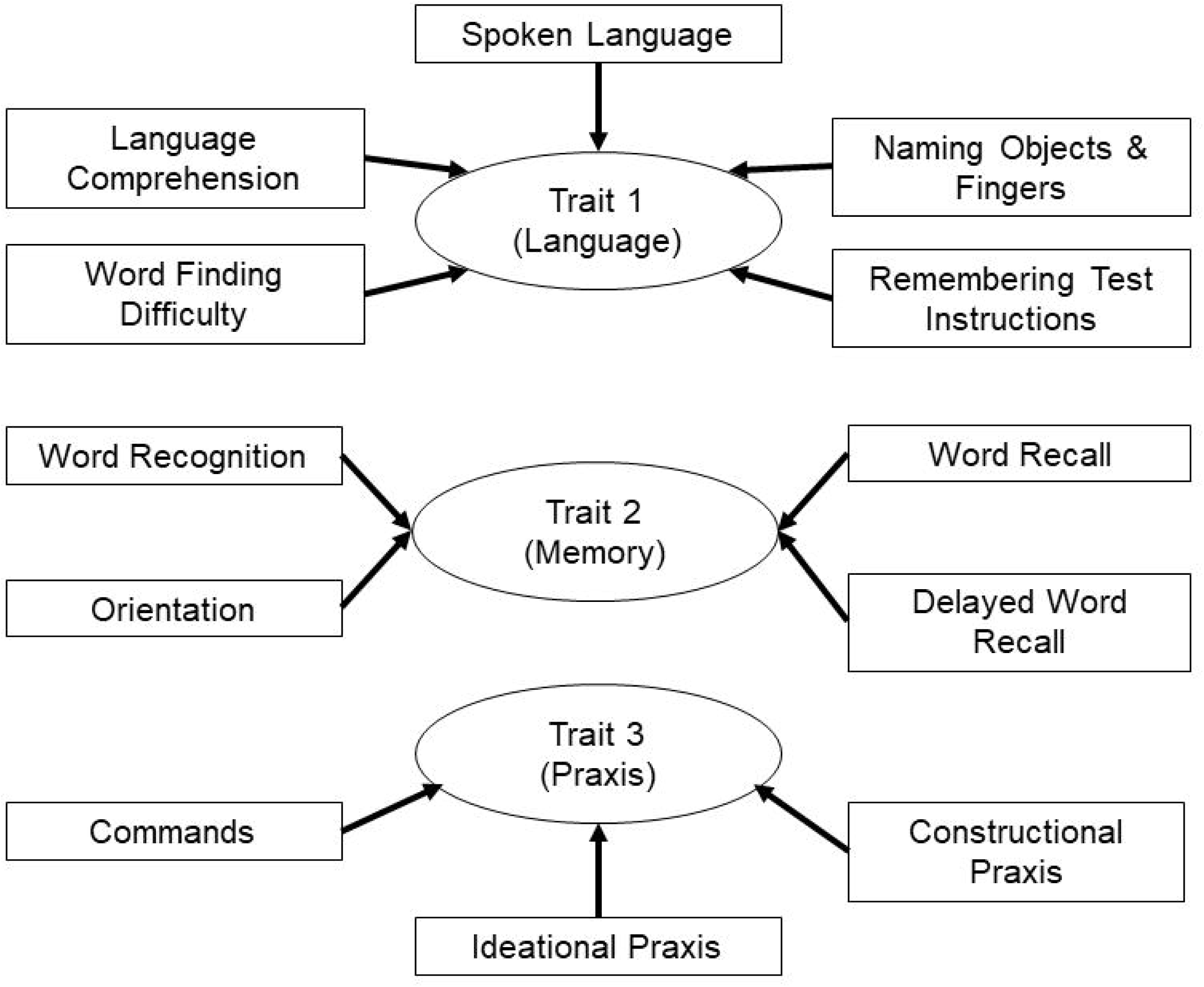
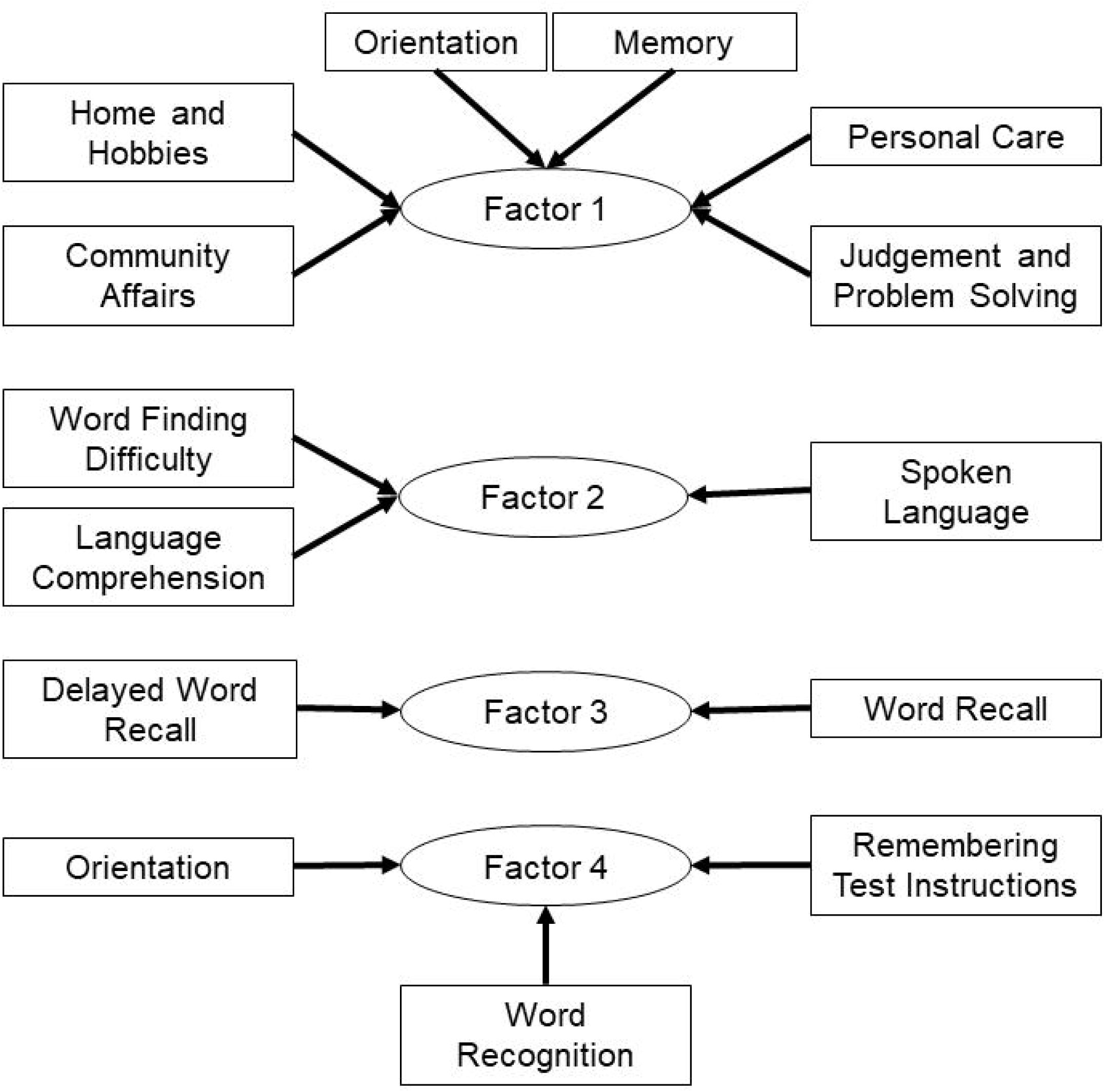
Grouping of ADAS-cog and CDR-sb sub-scales. (2A) Sub-scales of the ADAS-cog 12 test were grouped based on traits for memory, language and praxis to account for the topography of tissue loss in AD depending on the stage of disease. (2B) Sub-scales of the ADAS-cog 12 and the CDR-sb were analyzed by PCA, which resulted in grouping of sub-scales into four factors that explained most of the variance in the dataset. Composite variables were then generated, which included sub-scales identified in each factor by PCA, and were named factors 1 through 4. Note, factor 1 also contains the orientation sub-scale from the ADAS-Cog in addition to the ones from the CDR-sb.

### Cerebrospinal fluid biomarker measurements

Cerebrospinal fluid samples from both before and after treatment were available for 55 subjects and CSF collection was performed using a standardized sub-study protocol described elsewhere (27). Briefly, CSF collection by lumbar puncture was carried out between the screening and the baseline visit (within the 21 day window) and at the treatment termination (between 78 and 82 weeks). The lumbar puncture was performed using routine antiseptic cleansing and anesthesia with the patient in a reclining/sitting position. The lumbar puncture was performed using a Spinal Needle Quincke Type Point 0.7×75 mm (75–90 mm) that was inserted between L3/L4 or L4/L5 interspaces. Approximately 10ml of CSF was collected in 15ml polypropylene tubes. After gentle mixing, samples were centrifuged at 2000 × g for 10 min at 4°C to remove cells and debris, and then 1ml aliquots were prepared using polypropylene cryovials and subsequently frozen at −80°C until further use. Levels of Aβ38, Aβ40, and Aβ42 were quantified using the Meso-scale discovery (MSD) platform as previously described (28). Levels of total tau and phosphorylated tau (P181) were measured using commercially available sandwich ELISA kits (INNOTEST; Fujirebio) as per the manufacturer’s instructions and as previously described (28). All analyses were performed by board-certified laboratory technicians blinded to clinical information. We applied CSF biomarker criteria using total tau (> 350 pg/ml) and P181 tau (> 60 pg/ml) cut-offs as defined by Hansson et al., 2006 (29). An Aβ42/Aβ40 ratio cut-off of < 0.82 was used based on the concordance figures with amyloid PET imaging (Blennow, unpublished data) to compare baseline values with the clinical AD diagnosis. In this subsample, 91% of clinically diagnosed AD subjects also met CSF biomarker criteria for AD.

### APOE genotyping

The Gentra Puregene Kit (Gentra Systems) was used to purify DNA from frozen whole blood according to the manufacturer’s instructions and as previously described. The EzWay Direct APOE Genotyping Kit, (Koma Biotechnology), was used in accordance with the manufacturer’s instructions; specifically amplified DNA fragments corresponding to different APOE alleles were separated by electrophoresis in a 3% Ethidium Bromide stained metaphor agarose gel. All genotypes were then verified using rapid PCR with high-resolution melting analysis according to the manufacturer’s instructions (Novallele Genotyping). Apolipoprotein E genotypes were available on a subset of subjects (n = 328).

### Statistical analyses

General demographic characteristics across AD subgroups by treatment within the modified intent to treat (mITT) dataset were compared using either ANOVA or the Chi-square test, as applicable. Mixed linear model (MLM) regression was used to examine the main effects and the interactions between treatment, time (time points of study visits at 13 weeks, 52 weeks and 78 weeks) and AD severity at baseline. As we were interested in the independent contributions of the baseline AD severity and treatment effect over time on the cognitive outcomes, these analyses were also adjusted to account for the effect modification by gender and confounding by APOE (coded as those with the presence of ε4, without ε4 and those with no genotype information since not all mITT subjects had APOE genotypes available) and the confounding effects of age at which subjects left education (referred to as “education” hereon). To account for the treatment effect modification observed in the subgroup analyses, this model also included interactions between time and APOE; treatment and APOE; time, treatment and APOE; time and gender; treatment and gender; and time, treatment and gender. Interactive terms were also included for education and time and for treatment and education to account for education imbalance across AD severity subgroups. All of these variables were considered fixed factors. Subjects and country were treated as random factors. The autoregressive covariance structure was used in these MLM analyses. The outcome variables included change in the total ADAS-Cog 12 scores and change in composite scores from the ADAS-Cog 12 for different cognitive traits, CDR-sb and the ADCOMS. We also applied Principal Component Analysis (PCA) to minimize multicollinearity and achieve dimension-reduction for data on sub-scales from the ADAS-Cog 12 and CDR-sb for all visits. This method was used as an unsupervised procedure for achieving data-reduction and for identifying treatment responses in subgroups of subjects based on their baseline AD severity. The Kaiser-Meyer-Olkin (KMO) measure of sampling adequacy and Bartlett’s test for sphericity were used to ensure adequacy for PCA analysis (KMO value of > 0.6 and Bartlett p value < 0.05). Variables with eigenvalues of ≥1 were retained and PCA components (Factors) were extracted using varimax with Kaiser normalization for rotation in order to simplify and clarify the data structure. Individual ADAS-Cog 12 and CDR-sb sub-scales having a correlation of > 0.4 within each PCA factor were then grouped according to their association with a specific factor identified by PCA and then labeled as factors 1 through 4 (Figure 2B). These composite variables were used as the outcome measures for further analysis by MLM as described above. *Post-hoc* stratification was performed if the interaction terms for treatment, time and baseline AD severity showed a p value ≤ 0.05. Due to the confounding by APOE genotypes or effect modification by gender, their influences were also examined by further stratifying the subjects by APOE genotype and baseline AD severity, and by gender and baseline AD severity. Changes in CSF levels of Aβ38, Aβ40, Aβ42, total tau and P181 tau were calculated by subtracting values of the samples collected at the final visit from the baseline visit for each subject. Given the small sample size for the CSF subset, group comparisons using ANOVA were limited to mild (MMSE ≥ 20) and moderate (MMSE < 20) AD severity categories only. P values ≤ 0.05 were considered significant and all analyses were conducted using SPSS version 24 (IBM, NY).

## Results

### General characteristics of the study population stratified by AD severity

Demographic characteristics of the AD subgroups stratified by treatment are presented in Table 1. There were no differences across AD subgroups for age, weight or for the time elapsed since the diagnosis of AD. Similarly, there were no differences in ethnicity and the presence of the APOE ε4 allele (p > 0.05). However, there were significant differences in education across AD subgroups by treatment. There were also significant differences in gender, with females being overrepresented in the nilvadipine-treated moderate AD group compared to all other groups. Mild AD subjects having MMSE ≥ 20 were further stratified using a 1-point incremental increase in the MMSE cut-off (Figure 3). This showed that very mild AD subjects, who had MMSE ≥ 25, benefited the most after nilvadipine treatment when compared to all other MMSE subgroups, as indicated by the smallest change from baseline on the ADAS-Cog 12 seen in this cut-off compared to others. We therefore stratified the study population as described above to allow us to focus on very mild AD subjects along with mild and moderate AD subjects (MMSE = 20-24 and MMSE ≤19, respectively).

|  | **Moderate AD** |  | **Mild AD** |  | **Very mild AD** |  |
| --- | --- | --- | --- | --- | --- | --- |
|  | *MMSE ≤ 19* |  | *MMSE 20-24* |  | *MMSE ≥ 25* |  |
|  | **Nilvadipine** | **Placebo** | **Nilvadipine** | **Placebo** | **Nilvadipine** | **Placebo** |
|  | N = 92 | N = 94 | N = 118 | N = 113 | N = 36 | N = 44 |
| **Age at Randomisation** | 71.80 (0.95) | 71.79 (0.85) | 74.57 (0.77) | 73.16 (0.71) | 72.38 (1.25) | 73.80 (1.16) |
| **Baseline MMSE** | 16.28 (0.24) | 16.23 (0.23) | 21.79 (0.13) | 22.16 (0.13) | 25.56 (0.09) | 25.39 (0.08) |
| **Baseline ADAS-Cog** | 43.04 (1.01) | 42.95 (1.03) | 30.74 (0.63) | 30.96 (0.69) | 24.53 (0.95) | 25.73 (1.07) |
| **Baseline CDR** | 7.13 (0.31) | 6.83 (0.27) | 4.54 (0.17) | 4.53 (0.22) | 3.14 (0.33) | 3.24 (0.28) |
| **Baseline ADCOMS** | 4.08 (0.11) | 4.02 (.11) | 2.79 (0.07) | 2.84 (0.09) | 2.04 (0.14) | 2.05 (0.10) |
| **Age left Education\*** | 14.82 (0.36) | 16.05 (0.43) | 16.62 (0.36) | 16.41 (0.35) | 18.61 (0.88) | 17.70 (0.65) |
| **Years since AD Symptoms** | 4.62 (0.28) | 4.56 (0.27) | 4.31 (0.24) | 4.36 (0.28) | 3.53 (0.27) | 3.49 (0.31) |
| **Years since AD diagnosis** | 2.04 (0.20) | 1.80 (0.18) | 1.62 (0.14) | 1.75 (0.18) | 1.33 (0.22) | 1.35 (0.22) |
| **Female N (%)** | 66 (71.7) | 57 (60.6) | 77 (65.3) | 69 (61.1) | 17 (47.2) | 21 (47.7) |
| **Caucasian N (%)** | 89 (96.7) | 91 (96.8) | 115 (97.5) | 110 (97.3) | 36 (100) | 43 (97.7) |
| **APOE4 Carrier\* N (%)** | 32/62 (51.6) | 33/64 (51.6) | 47/75 (62.7) | 48/79 (60.8) | 15/24 (62.5) | 19/25 (76.0) |
| **Height at Baseline (cm)** | 162.4 (1.07) | 164.1 (0.91) | 163.7 (0.87) | 164.8 (0.81) | 165.3 (1.98) | 166.2 (1.43) |
| **Weight at Baseline (kg)** | 67.2 (1.19) | 71.0 (1.49) | 66.9 (1.09) | 69.5 (1.33) | 69.8 (2.31) | 68.5 (1.99) |
| **BMI at Baseline** | 25.5 (0.42) | 26.4 (0.50) | 25.0 (0.36) | 25.6 (0.42) | 25.5 (0.65) | 24.7 (0.49) |

\*Note: APOE genotyping was available only for a subset of individuals and the age the subjects left education was significantly different across MMSE categories. $P < 0.05$ . Education information was not available for 6 subjects and time since AD diagnosis was unavailable for 1 subject. Although the mITT dataset was composed of 498 subjects, 1 subject withdrew consent and therefore data on 497 subjects were available for analysis.

**Figure 3:**
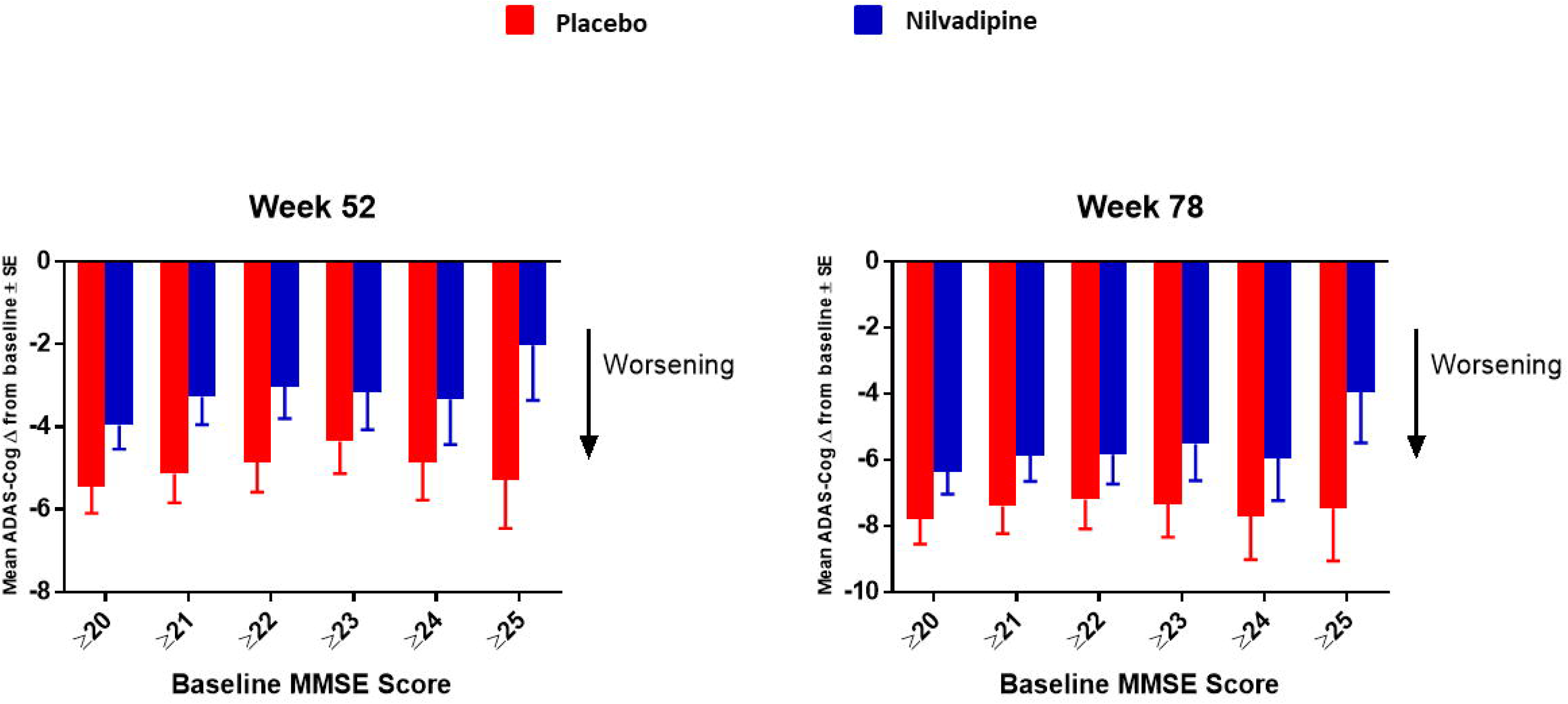
Further stratification of the mild AD group by increasing the increment of MMSE scores by 1 starting from ≥ 20 up to ≥ 25. Mean ± SE (for MMSE ≥ 20 n = 154 for nilvadipine and n = 157 for placebo; ≥ 21 n = 125 for nilvadipine and 136 for placebo; ≥ 22 n = 103 for nilvadipine and n = 117 for placebo; ≥ 23 n = 70 for nilvadipine and n = 99 for placebo; ≥ 24 n = 57 for nilvadipine and n = 68 for placebo; ≥ 25 n = 36 for nilvadipine and n = 44 for placebo). Mean change from baseline for the total ADAS-Cog 12 scores show the least decline among nilvadipine-treated subjects compared to placebo-treated subjects with MMSE score of ≥ 25.

### Changes in the ADAS-Cog 12 scores and the ADCOMS in response to nilvadipine treatment are modified by the baseline severity of AD

Regression analyses showed a significant interaction between nilvadipine treatment, baseline AD severity and time (F = 2.56, p = 0.02, Figure 4) on the change in the ADAS-Cog 12 after adjusting for the confounding factors described above. *Post-hoc* stratifications of the mean changes on the ADAS-Cog 12 showed a trend for (p = 0.06 at 52 weeks) less decline after nilvadipine treatment among very mild AD subjects and a greater decline on the ADAS-Cog 12 among nilvadipine-treated individuals with moderate AD (p = 0.02). There were no differences on the total ADAS-Cog 12 scores between nilvadipine-and placebo-treated individuals from the mild AD group (p > 0.05). There was also an interaction between treatment, time and baseline AD severity for the change in the ADCOMS (F = 2.16, p = 0.046). *Post-hoc* stratification showed significantly less decline on the ADCOMS in very mild AD subjects (p = 0.04 at 78 weeks), no change in mild AD subjects and a greater decline in moderate AD subjects (p = 0.03 at 78 weeks) who were treated with nilvadipine compared to their respective placebo treated groups. There were no differences on the CDR-sb total score with respect to the disease severity and treatment (data not shown). Since previous subgroup analyses showed an influence of the APOE ε4 allele and gender on the treatment outcome, supplementary figure 1 shows further stratification of changes in ADAS-Cog 12 and the ADCOMS in very mild, mild and moderate AD subjects by APOE ε4 status and gender. Compared to placebo, nilvadipine-treated, very mild AD individuals perform better on the ADCOMS over the 18-month period irrespective of the ε4 status or gender.

**Figure 4:**
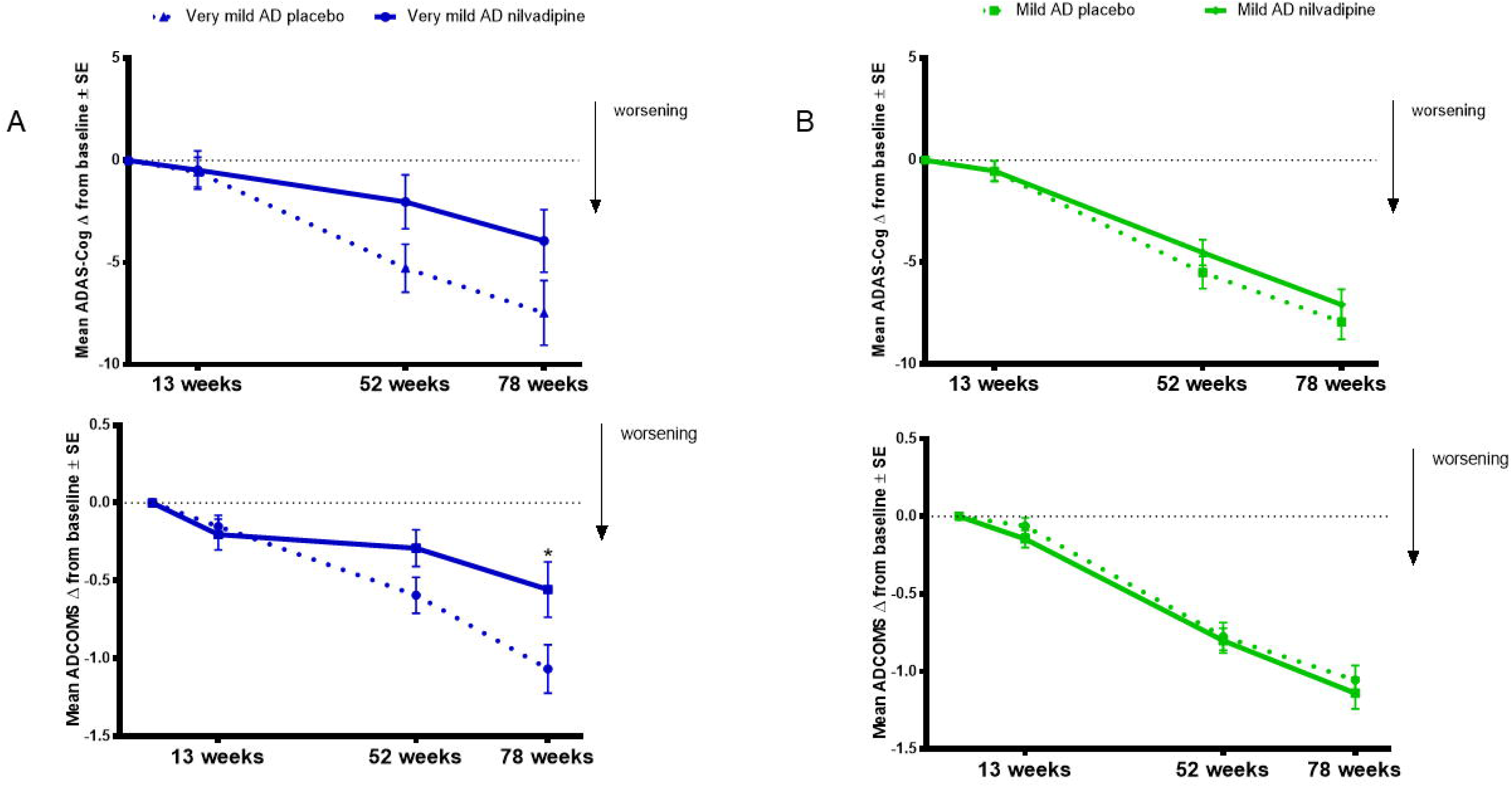
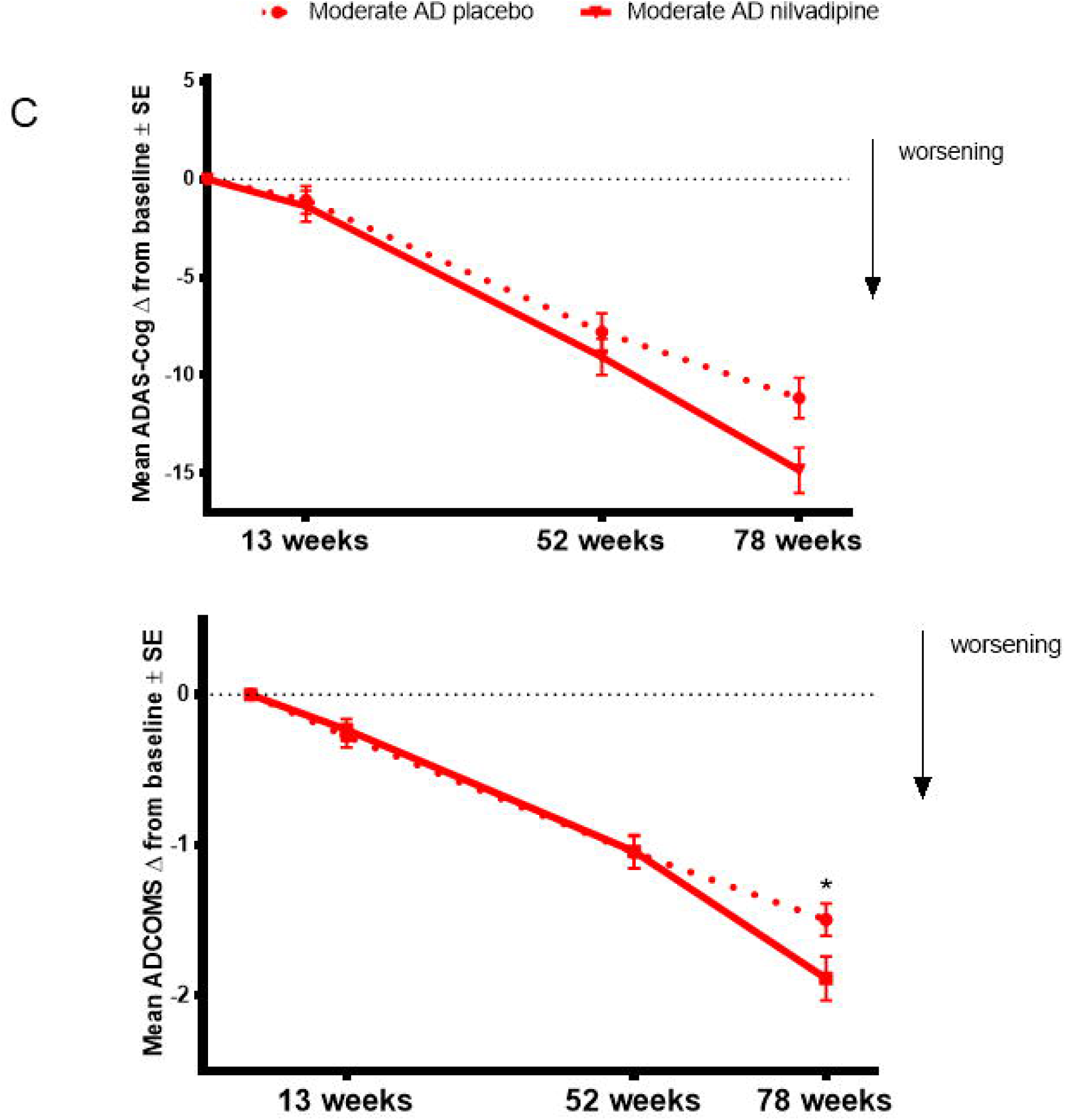
Nilvadipine-treated very mild AD subjects show less cognitive decline compared to controls on the ADAS-Cog 12 and the ADCOMS tests. Mean ± SE (n = 82 for moderate AD (MMSE ≤ 19) on nilvadipine, n = 94 for moderate AD on placebo, n = 118 for mild AD (MMSE 20-24) on nilvadipine, n = 113 for mild AD on placebo, n = 36 for very mild AD (MMSE ≥ 35) on nilvadipine and n = 44 for very mild AD on placebo) for the change in ADAS-Cog 12 scores. There was a significant effect for the interaction between treatment, time and baseline AD severity as assessed by MMSE scores after correcting for the confounding effects of APOE, gender and education, p < 0.05. (4A-B) Stratifications show that very mild AD subjects treated with nilvadipine have lower scores on the ADAS-Cog 12 and the ADCOMS compared to placebo after 78 weeks. Post-hoc analysis stratified by time show a significant treatment effect at 78 weeks for the ADCOMS. (4C-D) Mild AD subjects treated with nilvadipine scored similarly to their placebo controls on both the ADAS-Cog 12 and the ADCOMS (4E-F). However, moderate AD subjects treated with nilvadipine scored higher on both the ADAS-Cog 12 and the ADCOMS at 78 weeks compared to those on placebo. * p < 0.05

### Responses to nilvadipine on memory and language traits of the ADAS-Cog 12 depend on the baseline severity of AD

Regression analyses of the ADAS-Cog 12 sub-scales grouped as memory, language and praxis traits showed that the memory trait was significantly affected by the interaction between treatment, time and AD severity at baseline (F = 2.18, p = 0.04, Figure 5A) after adjusting for the confounders/effect modifiers as above. *Post-hoc* stratifications show that only very mild AD subjects treated with nilvadipine had less decline on the memory trait compared to placebo-treated very mild AD subjects (p = 0.04 at 52 weeks). There were no differences for the memory trait between nilvadipine-and placebo-treated mild AD subjects, while a non-significant decline on the memory trait was noted for moderate AD subjects treated with nilvadipine compared to placebo. Once adjusted for confounders, there was also a significant interaction between treatment, time and baseline AD severity on the language trait (F = 2.1, p = 0.05, Figure 5B), with *post-hoc* stratifications showing less decline in the language trait for the nilvadipine-treated mild AD group only (p = 0.03 at 52 weeks). There were no effects by AD severity, time or treatment on the praxis trait (p > 0.05).

**Figure 5:**
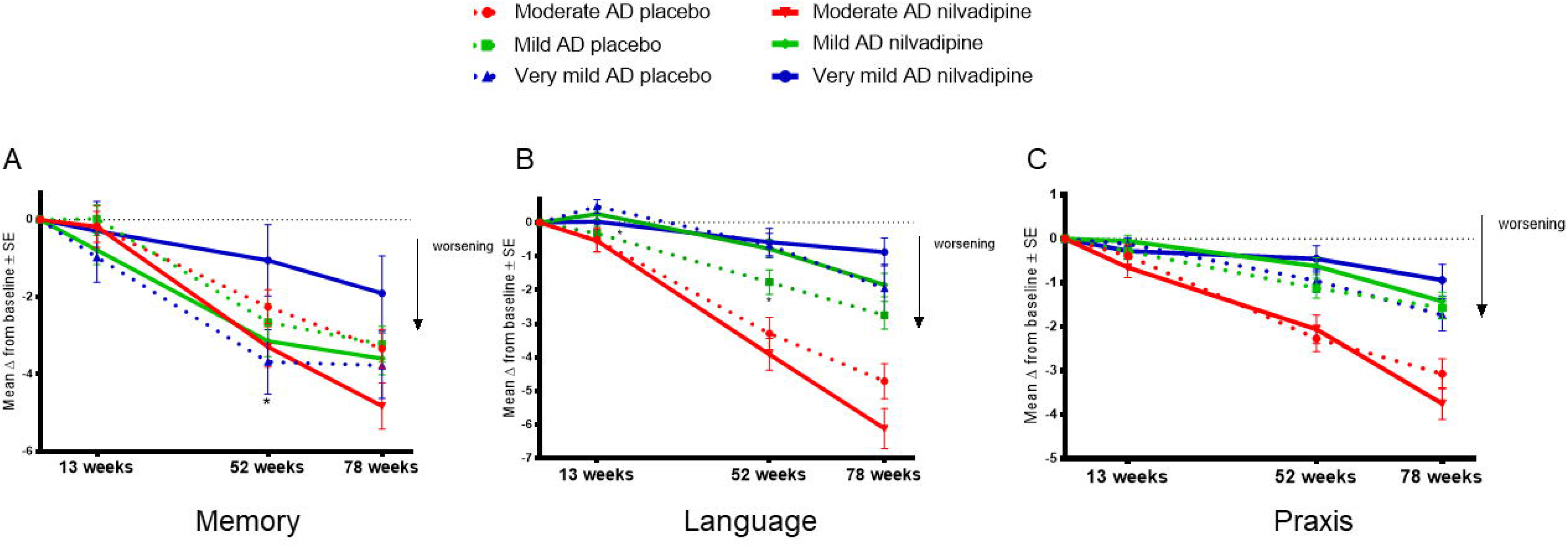
Very mild AD subjects show less decline on the memory trait, whereas mild AD subjects show less decline on the language trait, after nilvadipine treatment compared to placebo. Mean ± SE (n = 82 for moderate AD on nilvadipine, n = 94 for moderate AD on placebo, n = 118 for mild AD on nilvadipine, n = 113 for mild AD on placebo, n = 36 for very mild AD on nilvadipine, n = 44 for very mild AD on placebo) for the change in memory, language and praxis traits of grouped ADAS-cog 12 sub-scales. (5A) There was a significant effect for the interaction between treatment, time and baseline AD severity on the memory trait. Post-hoc stratifications by time show that very mild AD subjects treated with nilvadipine had significantly less decline on the memory trait compared to their controls. (5B) There was also a significant interaction between treatment, time and baseline AD severity for the language trait. Post-hoc stratifications by time show that mild AD subjects treated with nilvadipine had less decline on the language trait compared to the placebo-treated mild AD subjects. (5C) There was no effect seen for the praxis trait. *p < 0.05.

### Principal component analysis of the ADAS-Cog 12 sub-scales show benefits of nilvadipine on memory in subjects with very mild AD and language in mild AD

The PCA meeting the KMO-Bartlett’s test criteria of sampling adequacy and data sphericity identified 4 factors. Supplementary table 1 shows correlations between these factors and the different ADAS-Cog 12 and CDR-sb sub-scales. Among the 4 factors identified by PCA, there was a trend for an interaction between time, treatment and AD severity for factor 2 (F = 1.98, p = 0.07) loaded with the ADAS-Cog 12 sub-scales related to spoken language ability, comprehension, and word finding difficulty in spontaneous speech. *Post-hoc* stratifications of the mean changes showed that factor 2 differed by treatment only in the mild AD group, see Figure 6. There was also an interaction between treatment, time and baseline AD severity for factor 3 (F = 3.72, p = 0.001) loaded with the ADAS-Cog 12 sub-scales for the immediate and delayed word recall tasks. *Post-hoc* stratification showed that factor 3 differed by treatment only in the very mild AD group (See Figure 6). There were no significant differences between the groups for factor 1, which was loaded with all the sub-scales of CDR-sb along with several ADAS-Cog 12 sub-scales, or by factor 4, which was loaded with the ADAS-Cog 12 sub-scales for orientation, word recognition task and remembering the test instructions.

**Figure 6:**
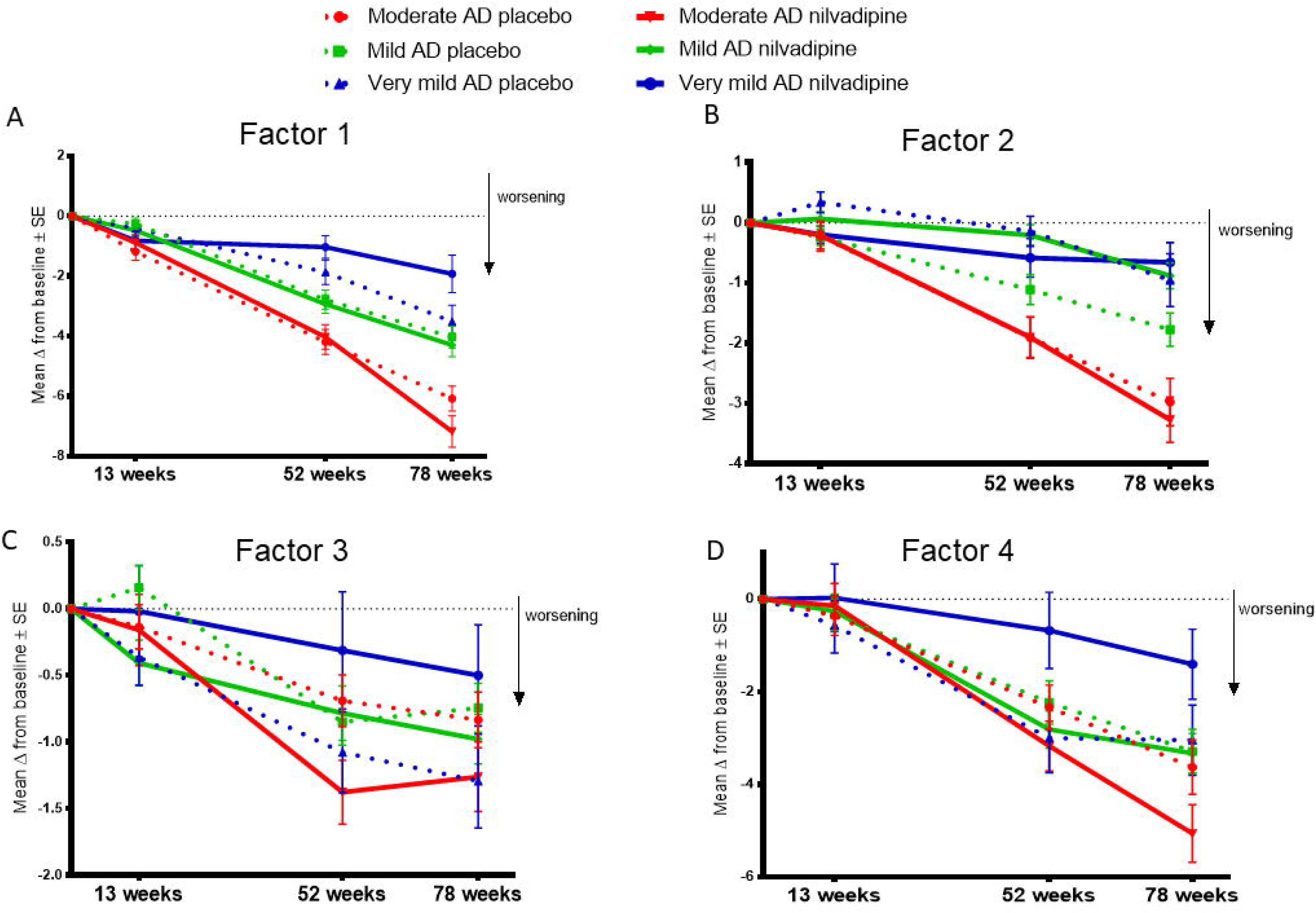
Nilvadipine-treated mild and very mild AD groups show less decline for PCA Factors 2 and 3, respectively. Mean ± SE (n = 82 for moderate AD on nilvadipine, n = 94 for moderate AD on placebo, n = 118 for mild AD on nilvadipine, n = 113 for mild AD on placebo, n = 36 for very mild AD on nilvadipine, n = 44 for very mild AD on placebo) for the change in Factors 1, 2, 3, and 4. (6A) There were no differences between any of the groups for Factor 1. (6B) A marginally significant interaction between time, treatment and AD severity was observed for Factor 2, p = 0.07, and subsequent stratifications show that only mild AD subjects treated with nilvadipine had less decline compared to their placebo controls. (6C) There was a significant interaction between time, treatment and AD severity for Factor 3, p < 0.05. (4D) There were no significant differences seen between groups for Factor 4.

### Nilvadipine treatment differentially modulates cerebrospinal fluid biomarkers depending on AD severity

Baseline demographics of the CSF subcohort stratified by mild and moderate AD status are provided in supplementary table 2. Changes in CSF Aβ42/Aβ40 ratios were significantly different across nilvadipine-and placebo-treated mild and moderate AD subjects (F = 3.55, p = 0.02, Figure 7A). *Post-hoc* analyses showed that CSF Aβ42/Aβ40 ratios showed a significant reduction in moderate AD subjects treated with nilvadipine compared to the placebo control group (p < 0.05). A trend for an increase in CSF Aβ42/Aβ40 ratios was observed in mild AD cases treated with nilvadipine compared to placebo (p = 0.067). Figures 7 and 8 show group differences between nilvadipine-and placebo-treated mild and moderate AD subjects for CSF Aβ38 (F = 2.98, p = 0.04), total tau (F = 6.29, p < 0.01) and P181 tau (F = 4.30, p < 0.01). *Post-hoc* analyses showed that in the moderate AD group, as compared to placebo, nilvadipine-treated subjects had significant increases in CSF Aβ38, total tau and P181 tau after nilvadipine treatment (p < 0.05). There was a non-significant increase in Aβ42 in nilvadipine-compared to placebo-treated mild AD subjects.

**Figure 7:**
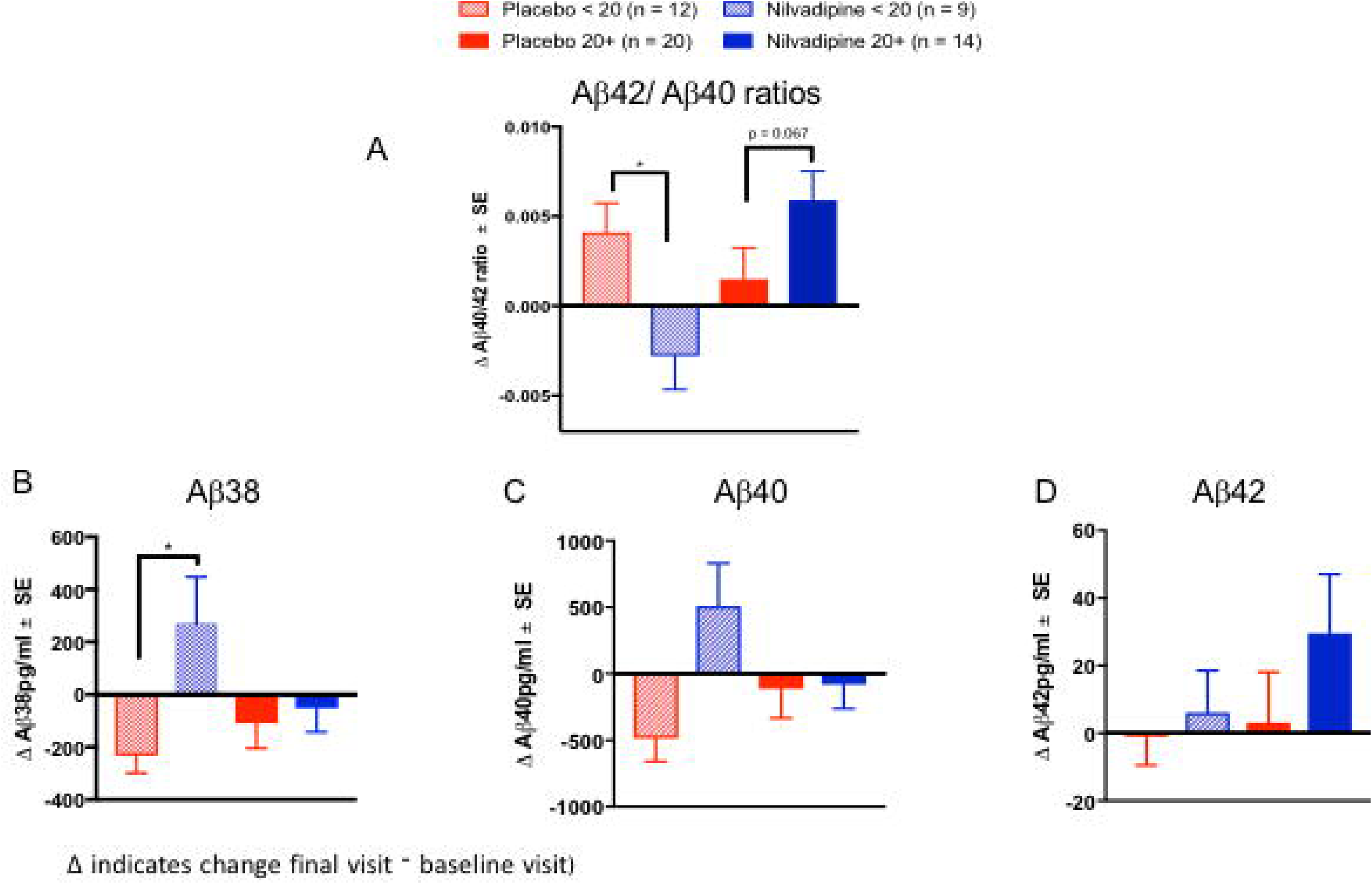
Ratios of CSF Aβ42/Aβ40 increase in nilvadipine-treated mild AD but decrease in moderate AD patients compared to their respective placebo groups. (Mean ± SE n = 9 for moderate AD on nilvadipine, n = 12 for moderate AD on placebo, n = 14 for mild AD on nilvadipine, n = 20 for mild AD on placebo). (A) Ratios of Aβ42/Aβ40 were higher in mild AD treated with nilvadipine compared to those treated with placebo (p = 0.067). There was a significant decrease in Aβ42/Aβ40 in moderate AD treated with nilvadipine compared to placebo. Also in moderate AD subjects, levels of (B) Aβ38 and (C) Aβ40 were elevated and (D) Aβ42 levels were unchanged in moderate AD subjects treated with nilvadipine. Levels of Aβ42 were non-significantly higher in mild AD treated with nilvadipine. *p < 0.05

**Figure 8:**
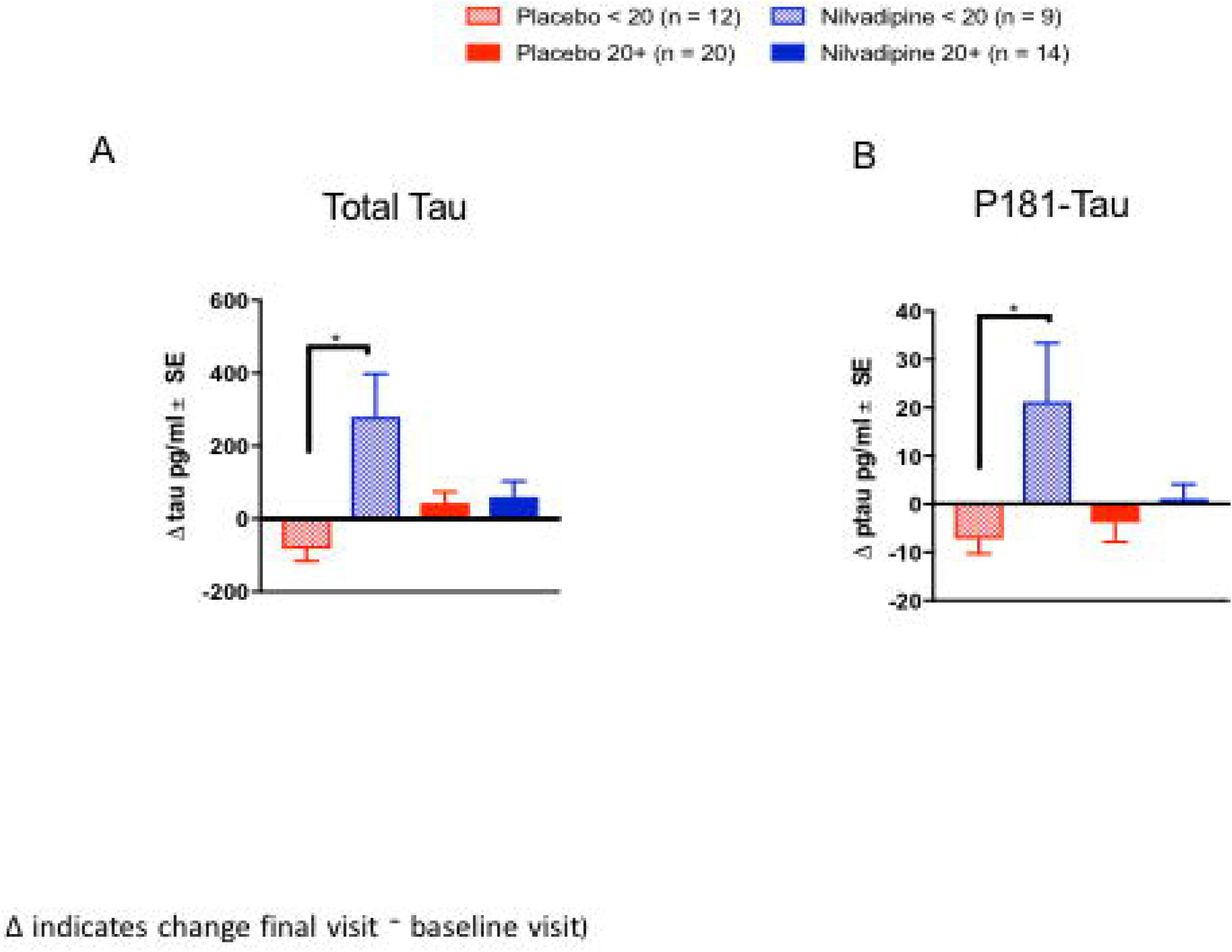
Total tau and P181Tau levels were increased in nilvadipine-treated moderate AD patients compared to their respective placebo groups. (Mean ± SE n = 9 for moderate AD on nilvadipine, n = 12 for moderate AD on placebo, n = 14 for mild AD on nilvadipine, n = 20 for mild AD on placebo). Levels of (A) total tau and (B) P181 tau were increased in moderate AD subjects treated with nilvadipine. In mild AD subjects, total tau or P181 tau did not differ between nilvadipine-treated and placebo control groups.*p < 0.05.

## Discussion

Many clinical trials in combined populations of mild and moderate AD patients have failed to show benefits, possibly because different stages of AD reflect changes in the extent of the underlying amyloid and tau pathologies. Frequently the same drugs that have failed in combined mild and moderate populations have shown suggestive benefits for subjects in early stages of the disease (7-10, 30), when the extent of amyloid and tau pathologies is considerably lower than for subjects in the moderate disease stage (31-33). Although analysis of the overall population showed no benefit of nilvadipine, our exploratory studies show that very mild AD subjects experienced less cognitive decline on the total ADAS-Cog 12 in the nilvadipine-treated group compared to placebo. However, moderate AD subjects treated with nilvadipine showed a greater decline compared to their controls. We also used the ADCOMS, which showed that the beneficial effects of nilvadipine were restricted to the very mild AD group. An examination of different cognitive traits from the ADAS-Cog 12 sub-scales showed that nilvadipine had beneficial effects on the memory trait in the very mild AD group and the language domain in the mild AD group. However, moderate AD patients showed worsening of the memory trait after nilvadipine intervention. These findings suggest a differential impact of nilvadipine depending on the stage of the disease at which treatment is initiated.

The examination of changes over time in the total ADAS-Cog 12 scores, stratified by different stages of AD, suggested that, after 78 weeks of treatment, very mild AD patients treated with nilvadipine showed less cognitive impairment than placebo controls. There were no differences in the total ADAS-Cog 12 scores between the placebo-and nilvadipine-treated subjects in the mild AD group, and the scores of the moderate AD subjects treated with nilvadipine showed greater cognitive decline compared to placebo. Placebo groups for very mild and mild AD declined at a similar rate, whereas the moderate AD placebo group declined at nearly twice the rate of the milder groups during this period. These findings are largely consistent with those previously reported for placebo groups in other clinical trials, in which a lower decline is generally noted in mild AD versus moderate AD subjects (34, 35). In any of the AD severity categories, there was no correlation between blood pressure changes and total ADAS-Cog 12 changes among those treated with nilvadipine (*pers. comm*). This is consistent with our preclinical studies showing that the anti-Aβ effects of nilvadipine are not related to its cerebrovascular effects (17).

We did not observe an effect of nilvadipine treatment on the changes in CDR-sb. Studies have shown that while the ADAS-Cog 12 test is useful at estimating progression in mild stages of AD, the CDR-sb test is a global impression scale designed for staging of dementia rather than quantifying cognitive change over time. It is therefore possible that the CDR-sb lacks the desired sensitivity to detect subtle cognitive changes due to high test-retest variability for detecting cognitive differences (36, 37). Since disease progression in AD differs by the initial stage of the disease, it has been argued that the ADAS-Cog 12 and CDR-sb tests alone do not have the desired sensitivity to detect subtle changes in cognitive decline that occur in mild AD subjects. Recently, Wang and colleagues developed a composite variable, ADCOMS, which uses subscales from the ADAS-Cog 12, the MMSE and the CDR-sb and assigns adjusted weights generated from PLS regression to identify their relative contributions to AD progression (26). This composite outcome includes both cognitive and functional measures. Many clinical trials now incorporate the ADCOMS as it seems to be sensitive at detecting treatment effects in the early stages of AD(38, 39). When we used the ADCOMS (modified to accommodate the absence of MMSE sub-scales), we observed cognitive benefits in very mild AD subjects treated with nilvadipine.

Subjects in different stages of AD demonstrate differential decline in memory, language and praxis traits. These traits can be mapped to the underlying brain tissue loss in AD in different stages of the disease. For instance, neuronal loss in early AD starts within the medial temporal lobe (MTL), which is primarily involved with memory function. With the advancement of AD, further degeneration occurs within the parietal, frontal, and occipital lobes, which are involved in language processing and praxis (24, 40). Our exploratory analyses of these cognitive traits suggest that benefits of nilvadipine were restricted to the memory trait in very mild AD subjects. In mild AD cases, who are at a more advanced disease stage compared to very mild cases, we observed a reduced decline for the language trait in nilvadipine-treated subjects. There were no effects of nilvadipine on praxis for any of the AD subpopulations. In the moderate AD group, there was no specific domain accounting for the overall decrement in ADAS-Cog 12 with nilvadipine treatment, but rather there were trends for decline in all cognitive domains. Over the 18 months, placebo-treated very mild AD subjects showed significant decline in memory. This is to be expected since very mild AD subjects initially have the most functional memory, which is rapidly lost in the early stages of the disease. The language trait remained largely preserved in very mild AD subjects treated with placebo, but continued to decline further in mild and moderate AD placebo groups. The praxis trait further declined in moderate AD on placebo with minimal decline in both very mild and mild AD on placebo. This is again to be expected as loss of praxis generally occurs after the loss of memory function as AD progresses. Collectively, these data suggest that examination of appropriate cognitive domains relevant to the stage of AD might improve our ability to evaluate treatment effects in AD clinical trials.

Alzheimer’s Disease patients in the early stages of the disease have limited cognitive and functional impairment. As such, currently available clinical outcome scales, such as those utilized here, may not be able to capture subtle cognitive changes, particularly in the presence of relatively small functional decline (26, 41, 42). As such, there remains a need for the development of composite scores for assessing treatment responses. One such approach includes using the ADCOMS (26), which was recently utilized in other AD trials as an outcome measure in 18-month trials to assess disease modification by treatment(39). We also used the ADCOMS and found that nilvadipine treatment was beneficial for very mild AD. Using another similar approach, we performed PCA on sub-scales from both the ADAS-Cog 12 and CDR-sb as an unsupervised dimension reduction method to group specific cognitive domains. Factor 1 loading primarily contained all of the CDR-sb sub-scales and also the orientation sub-scale from ADAS-Cog 12, and remained unaffected by nilvadipine treatment. Factor 2 loading contained language and word finding abilities-related sub-scales from the ADAS-Cog 12 and, as with the trait analysis, a trend for potential benefit of nilvadipine was limited to the mild AD group only. Factor 3 was loaded with immediate and delayed recall, and showed less decline after treatment in the very mild AD group. Factor 4 loading contained orientation, word recognition and remembering test instructions, which did not differ between the nilvadipine and placebo groups in any of the AD severity subpopulations. Collectively, these results suggest that memory was preserved in very mild AD and language was preserved in mild AD subjects after nilvadipine treatment.

We also stratified the study population by AD severity at baseline and by APOE ε4 carrier status, and our results suggested that among non-carriers, very mild AD cases had lower decline with nilvadipine treatment, but AD patients in the moderate stage showed cognitive decline with nilvadipine treatment. In the subpopulation that was genotyped for APOE, ε4 carriers were (unusually) as cognitively intact as non-carriers at baseline. Despite this, ε4 carriers did not benefit from nilvadipine treatment regardless of the initial disease stage, which is consistent with the literature from other AD treatment trials (43). However, the very mild and moderate ε4 carrying placebo groups did not decline between months 12 and 18. Stratification by gender showed that both males and females in the very mild AD stage had less cognitive decline with nilvadipine treatment compared to placebo treatment, while females in the moderate AD stage showed a greater cognitive decline with nilvadipine treatment. With respect to gender, moderate females did worse on nilvadipine than males. This may be consistent with some other studies which show poor response to certain therapies in AD females compared to AD males (44).

Correlative studies of amyloid imaging with CSF Aβ levels show that the decrease in CSF Aβ42 is an early event in AD pathogenesis (31). Longitudinal profiling of CSF Aβ in AD animal models shows that, with AD progression, CSF Aβ42 levels decline first but Aβ40 levels do not fall until the very late stages of AD pathology (45). In addition, recent clinical studies have shown that CSF Aβ42/Aβ40 ratios have a better concordance with amyloid PET imaging for biomarker-based diagnosis of AD than using either Aβ42 or Aβ40 alone (46), and that this ratio is consistently low in AD subjects with high brain amyloid deposition (46, 47). In the present study, in mild AD patients, CSF Aβ42/Aβ40 ratios increased in nilvadipine-treated subjects compared to placebo-treated subjects. The mild AD placebo group essentially had no change in Aβ40 and Aβ42 levels. Based on the known mechanism of action of nilvadipine (15), we speculate that the increase in CSF Aβ ratios in the mild AD nilvadipine-treated group reflects clearance of Aβ42 across biological barriers since there was a trend for an increase in Aβ42 levels in nilvadipine-treated subjects. However, moderate AD subjects treated with nilvadipine had a decline in CSF Aβ42/Aβ40 ratios compared to their placebo-treated controls. This was due to a minimal increase in Aβ42 relative to a larger increase in Aβ40 levels in this moderate AD group. Nevertheless, a decline in CSF Aβ42/Aβ40 corresponded with the worsening of cognition after nilvadipine treatment in this group. Based on preclinical nilvadipine studies (15), the observation of increased Aβ40 and Aβ438 in the moderate AD subjects treated with nilvadipine could be interpreted as increased clearance of these species, allowing shorter and more soluble Aβ species than Aβ42 to move out of the brain and into CSF. However, we do not yet understand the mechanism by which nilvadipine caused moderate AD subjects to experience a greater cognitive decline than placebo treated subjects. One explanation could be an increased production of Aβ species, which is probably unlikely since Aβ42 levels remained essentially unaltered in moderate AD subjects treated with nilvadipine. An increase in CSF Aβ42/Aβ40 ratios was observed in moderate AD subjects treated with placebo and appears to be a consequence of decline in Aβ40 without any further drop in Aβ42, which is to be expected in such advanced stages of AD (45). Total tau and P181 tau were increased after nilvadipine treatment in moderate AD subjects. Interestingly, placebo-treated mild and moderate AD subjects showed a decline in total tau and phospho-tau, which is unexpected, but has been previously reported in a longitudinal study of AD subjects (48). Together, biomarker data from this NILVAD trial suggest that cognitive improvement in mild AD subjects after treatment corresponds with an increase in CSF Aβ42/Aβ40 ratios, whereas worsening of cognition in moderate AD subjects is paralleled by a decrease in CSF Aβ42/Aβ40 ratios and higher total and phospho-tau levels.

## Conclusion

With failures of most AD trials to satisfy efficacy criteria in mixed AD populations, exploratory analyses of existing trial data are justified and necessary to understand lack of efficacy and to identify if sub-populations exist that may have benefited from interventions. The NILVAD trial was designed for the analysis of a mixed mild and moderate AD population and further stratification of the study population into very mild, mild and moderate AD was unplanned and therefore exploratory. As such, these subgroup analyses were underpowered, particularly when considering the confounding effects of gender and APOE. Nevertheless, analyses adjusted for these factors continue to suggest that very mild AD subjects responded positively to nilvadipine on both the ADAS-Cog 12 and the ADCOMS. Furthermore, analyses of the ADAS-Cog 12 sub-scales demonstrate that memory and language traits are beneficially impacted by nilvadipine treatment in very mild and mild AD patients, respectively. Together, findings from the clinical studies and CSF biomarker analyses suggest a differential response to nilvadipine treatment in AD based on the severity of the disease at treatment initiation. These findings are also consistent with the results of several other experimental AD treatments where only very early stage AD subjects demonstrated benefit, such as Solanezumab (7, 10, 30), aducanumab (8) and LipiDiDiet trials (9). Consequently, the Alzheimer’s therapeutic field is increasingly targeting the early stages of AD (49). Finally, possible benefits in the very mild AD group identified by these exploratory analyses warrant further studies of nilvadipine treatment in very mild, prodromal or even preclinical AD patients.

## Declarations

### Acknowledgments

The authors thank all participants in the NILVAD consortium as well as all of the patients and caregivers who so generously gave of their time. The authors also thank Muireann O’Briain; Oliver Gupta; the members of the Ethics Advisory Board (Ursula Collins, Mary Donnelly, Tony O’Brien and Shaun O’Keefe); the members of the Safety Advisory Board (Paul Aisen, Suzanne Hendrix, Robin Jacoby and Maurice O’Connell); and the members of the Data Safety Monitoring Board (Bernadette McGuinness, John Newell, Martin O’Donnell and Peter Passmore). The authors wish to thank all other study partners – Alzheimer Europe and Newsweaver who helped with dissemination and promotion of the trial and GABO:mi who were the project management company for the majority of the trial. We also would like to acknowledge LabeX DISTALZ for facilitating subject recruitment within the Hauts-de-France region.

### Ethics statement

The list of IRB by each country where the study approval was received:

France: Comite de protection des personnes nord quest III

Greece: Scientific Council of Papanikolaou Hospital Thessaloniki

Holland: Radboud universitair medisch centrum Concernstaf Kwaliteit en Veiligheid Commissie Mensgebonden Onderzoek Regio Arnhem-Nijmegen (Chair: M.J.J. Prick)

Hungary: Medical Research Council Ethics Committee for Clinical Pharmacology (KFEB) (Chair: Dr Fϋrst Zsuzsarina)

Italy: Comitato Etico Istituzioni Ospedaliere Cattoliche(Chair: Dr. Giovanni Zaninetta),

Comitato Etico IRCCS MultiMedica (Chair: Prof Emilio Trabucchi), Comitato Etico dell’Aienda Ospedaliera Universitaria S. Martino di Genova (Chair: Dott. Luigi Francesco Meloni) And Comitato Etico Regione Liguria (Chair: Prof. Fulvio Brema), Comitato Etico Fondazione Don Carlo Gnocchi (Chair: Prof Flaminio Cattabeni),

Sweden: Regionala etikprӧvningsnamnden I Gӧteborg (Chair: Vastra Gӧtalandsregionen)

United Kingdom: NRES Committee London – Harrow (Chair: Dr Jan Downer and Miss Shelly Glaister-Young)

Ireland: Tallaght Hospital / St. James’s Hospital Joint Research Ethics Committee (Chair: Dr Peter Lavin)

Germany: Ethik-Kommission der Bayerischen Landesarztekammer (Chair: Prof. Dr. med. Joerg Hasford)

### Consent for Publication

Not applicable

### Availability of Data and Material

All de-identified research data are available upon request.

### Authors Contributions

Conceptualization: Abdullah L, Crawford F, Mullan M and Lawlor B.

Data Curation: Abdullah L, Langlois H, Tsolaki M, Börjesson-Hanson A, Olde Rikkert M, Pasquier F, Wallin A, Kennelly S and Blennow K

Methodology: Abdullah L, Langlois L, and Hendrix S

Project administration and Supervision: Abdullah L, Crawford F, Lawlor B and Mullan M

Writing – review & editing: Abdullah L, Crawford F, Langlois H, Tsolaki M, Börjesson-Hanson A, Olde Rikkert M, Pasquier F, Wallin A, Kennelly S, Hendrix S, Blennow K, Lawlor B, Mullan M

## Abbreviations

AD: Alzheimer’s Disease
mITT: modified Intent-to-Treat
ADAS-Cog12: Alzheimer’s Disease Assessment Scale Cognitive 12
CDR-sb: Clinical Dementia Rating Scale sum of boxes
ADCOMS: AD composite score
CSF: Cerebral Spinal Fluid
DHP: Dihydropyridine
CCB: Calcium channel blocker
MCI: Mild Cognitive Impairment
BBB: blood brain barrier
Syk: spleen tyrosine kinase
IRB: Institutional Review Board
NINCDS-ADRDA: National Institute of Neurological and Communicative Disorders and Stroke/Alzheimer’s Disease and Related Disorders Association Inc.
MMSE: Mini-Mental State Examination
PLS: Partial Least Square
MSD: Meso Scale Discovery
PCA: Principal Component Analysis
KMO: Kaiser-Meyer-Olkin
MTL: medial temporal lobe

---

**Supplemental figure 1: Changes in total ADAS-Cog 12 stratified by baseline AD severity and APOE ε4 carrier status.** Mean ± SE (n = 32 for moderate AD on nilvadipine, n = 33 for moderate AD on placebo, n = 47 for mild AD on nilvadipine, n = 48 for mild AD on placebo, n = 15 for very mild AD on nilvadipine, n = 19 for very mild AD on placebo). Moderate non-carrier AD subjects treated with nilvadipine showed decline over the 78-week period, whereas mild or moderate ε4 carrier AD subjects treated with nilvadipine scored similarly to their placebo controls.

