## Supplementary Table 1 for "Exploratory analyses suggest less cognitive decline with nilvadipine treatment in very mild Alzheimer’s disease subjects"

|  | Factor 1 | Factor 2 | Factor 3 | Factor 4 |
| --- | --- | --- | --- | --- |
| Memory* | 0.665 |  |  |  |
| Orientation* | 0.688 |  |  |  |
| Judgment and Problem Solving* | 0.677 |  |  |  |
| Community Affairs* | 0.705 |  |  |  |
| Home and Hobbies* | 0.720 |  |  |  |
| Personal Care* | 0.609 |  |  |  |
| Word Recall task |  |  | 0.779 |  |
| Naming Fingers and Objects |  |  |  |  |
| Commands |  |  |  |  |
| Constructional Praxis |  |  |  |  |
| Delayed Word Recall |  |  | 0.752 |  |
| Ideational Praxis |  |  |  |  |
| Orientation | 0.459 |  |  | 0.513 |
| Word Recognition Task |  |  |  | 0.671 |
| Spoken Language Ability |  | 0.822 |  |  |
| Comprehension |  | 0.767 |  |  |
| Word-finding difficulty in spontaneous speech |  | 0.759 |  |  |
| Remembering Test Instructions |  |  |  |  |

**Supplementary table 1:** Factors identified by PCA from ADAS-Cog 12 and CDR-sb subscales

Note: Extraction Method: Principal Component Analysis.  Rotation Method: Varimax with Kaiser Normalization.^a^ Values for each subscale within each factor represents correlation coefficient generated by PCA analysis.
