## Supplementary Table 2 for "Exploratory analyses suggest less cognitive decline with nilvadipine treatment in very mild Alzheimer’s disease subjects"

|  | Moderate AD | | Mild AD | |
| --- | --- | --- | --- | --- |
|  | *MMSE < 20* | | *MMSE ≥ 20* | |
|  | Nilvadipine | Placebo | Nilvadipine | Placebo |
|  | N = 9 | N = 12 | N = 14 | N = 20 |
| Age at Randomisation | 66.8 (2.8) | 67.6 (2.8) | 67.7 (2.4) | 68.3 (1.5) |
| Baseline MMSE | 16.3 (0.4) | 15.5 (0.8) | 22.9 (0.5) | 23.4 (0.5) |
| Baseline ADAS-Cog | 38.1 (2.2) | 44.6 (3.5) | 29.8 (2.3) | 30.4 (1.6) |
| Baseline CDR | 4.4 (0.4) | 5.7 (0.5) | 4.8 (0.5) | 4.3 (0.4) |
| Age left Education* | 16.8 (0.9) | 15.5 (1.2) | 18.9 (1.0) | 16.6 (0.8) |
| Years since AD Symptoms | 4.9 (0.8) | 4.4 (0.8) | 5.1 (0.9) | 3.8 (0.5) |
| Years since AD diagnosis | 1.4 (0.5) | 1.0 (0.4) | 1.5 (0.4) | 1.2 (0.3) |
| Female  N (%) | 7 (77.8) | 7 (58.3) | 9 (64.3) | 7 (35.0) |
| Caucasian  N (%) | 9 (100.0) | 12 (100.0) | 14 (100.0) | 20 (100.0) |
| APOE4 Carrier*  N (%) | 5 (55.6) | 4 (33.3) | 11 (78.6) | 13 (65.0) |
| Height at Baseline (cm) | 163.6 (3.6) | 165.3 (2.5) | 165.6 (2.2) | 168.8 (2.1) |
| Weight at Baseline (kg) | 74.4 (4.7) | 78.3 (2.9) | 64.0 (3.1) | 73.5 (2.5) |
| BMI at Baseline | 27.8 (1.5) | 28.8 (1.3) | 23.4 (1.2) | 25.8 (0.7) |
| Baseline Aß38 (pg/ml) | 2211.9 (213.7) | 2260.3 (244.8) | 2296.3 (151.0) | 2359.3 (170.6) |
| Baseline Aß40 (pg/ml) | 5231.9 (475.0) | 5460.5 (499.2) | 5475.2 (277.3) | 5818.0 (351.8) |
| Baseline Aß42 (pg/ml) | 292.5 (29.9) | 247.2 (24.8) | 284.0 (27.6) | 294.0 (21.4) |
| Baseline Total Tau (pg/ml) | 761.8 (118.4) | 877.3 (111.7) | 740.0 (47.5) | 776.7 (55.6) |
| Baseline P-Tau (pg/ml) | 71.8 (9.4) | 78.3 (9.2) | 71.6 (4.8) | 77.7 (5.4) |
| Baseline Aß42/Aß40 | 0.06 (0.005) | 0.05 (0.004) | 0.05 (0.004) | 0.05 (0.004) |

**Supplementary table 2:**

Note: gender differences were significantly different between placebo and nilvadipine treated individuals (p < 0.05).
