## Supplementary figures and images for "Exploratory analyses suggest less cognitive decline with nilvadipine treatment in very mild Alzheimer’s disease subjects"

### Supplemental Figure 1A

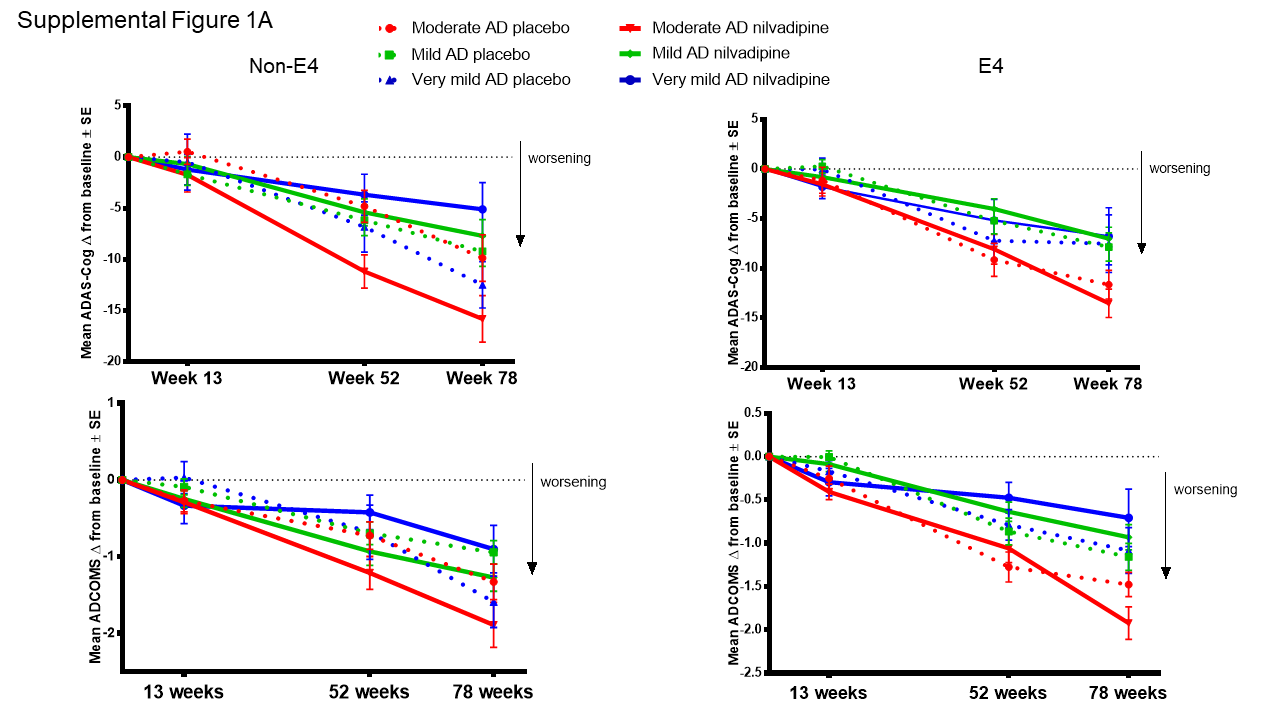

### Supplemental Figure 1B

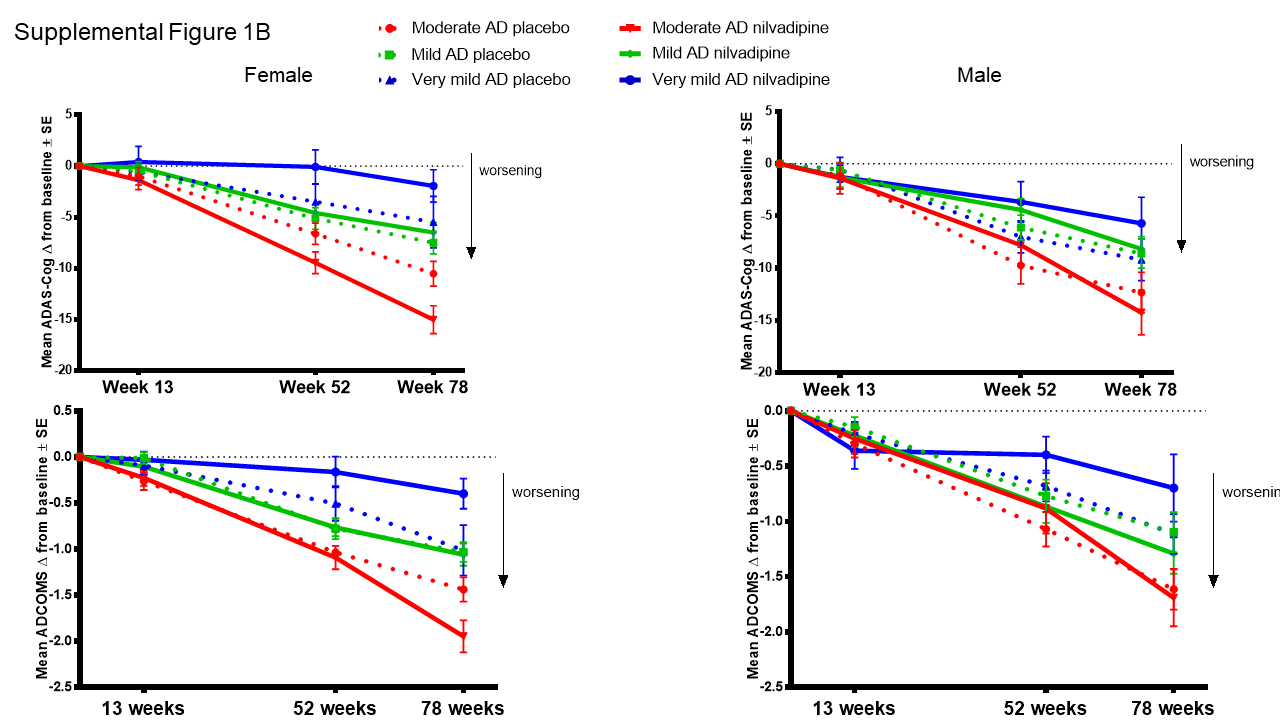
